# Dynamic control systems that mimic natural regulation of catabolic pathways enable rapid production of lignocellulose-derived bioproducts

**DOI:** 10.1101/2022.01.12.475730

**Authors:** Joshua R. Elmore, George Peabody, Ramesh K. Jha, Gara N. Dexter, Taraka Dale, Adam Guss

**Affiliations:** Oak Ridge National Laboratory, Biological and Environmental Systems Sciences Division, One Bethel Valley Road, Oak Ridge, TN 37830; Pacific Northwest National Laboratory, 902 Battle Blvd, Richland, WA 99352; Los Alamos National Laboratory, Biosciences Division, 1663 Bikini Atoll Road, Los Alamos, NM 87545

## Abstract

Expanding the catabolic repertoire of engineered microbial bioproduction hosts enables more complete use of complex feedstocks such as lignocellulosic hydrolysates and deconstructed mixed plastics, but the deleterious effects of existing expression systems limit the maximum carry capacity for heterologous catabolic pathways. Here, we demonstrate use of a conditionally beneficial oxidative xylose catabolic pathway to improve performance of a *Pseudomonas putida* strain that has been engineered for growth-coupled bioconversion of glucose into the valuable bioproduct *cis,cis*-muconic acid. In the presence of xylose, the pathway enhances growth rate, and therefore productivity, by >60%, but the metabolic burden of constitutive pathway expression reduces growth rate by >20% in the absence of xylose. To mitigate this growth defect, we develop a xylose biosensor based on the XylR transcription factor from *Caulobacter crescentus* NA1000 to autonomously regulate pathway expression. We generate a library of engineered xylose-responsive promoters that cover a three order-of-magnitude range of expression levels to tune pathway expression. Using structural modeling to guide mutations, we engineer XylR with two and three orders-of-magnitude reduced sensitivity to xylose and L-arabinose, respectively. A previously developed heterologous xylose isomerase pathway is placed under control of the biosensor, which improves the growth rate with xylose as a carbon source by 10% over the original constitutively expressed pathway. Finally, the oxidative xylose catabolic pathway is placed under control of the biosensor, enabling the bioproduction strain to maintain the increased growth rate in the presence of xylose, without the growth defect incurred from constitutive pathway expression in the absence of xylose. Utilizing biosensors to autonomously regulate conditionally beneficial catabolic pathways is generalizable approach that will be critical for engineering bioproduction hosts bacteria with the wide range of catabolic pathways required for bioconversion of complex feedstocks.

## INTRODUCTION

Efficient conversion of lignocellulose to sustainably produced transportation fuels and industrial chemicals will be critical for reducing dependence on petroleum^1, 2^, and achieving this goal is contingent upon improvements to the carbon efficiency of lignocellulosic feedstock bioconversion^1, 3-6^. In addition to abundant monosaccharides (e.g., glucose, xylose) lignocellulosic hydrolysates are composed of other monosaccharides, oligosaccharides, organic acids, inhibitory compounds, aromatics, proteins (from biomass and enzyme cocktails), and other diverse compounds^7^. These compounds comprise a substantial amount of carbon available for bioconversion, but no known organism can consume all of them. Accordingly, engineering microbial hosts with such expansive catabolic capabilities is a substantial metabolic engineering challenge that must be overcome for carbon efficient bioconversions. This is particularly important for production of biochemicals and other non-fuel products, where residual feedstock components must be removed in expensive separations steps^8, 9^. This challenge also provides an opportunity to employ innovative metabolic engineering strategies that are not feasible when using current homogenous industrial feedstocks (*e*.*g*. glucose, starches, vegetable oils). Engineering strategies that utilize heterologous pathways to funnel multiple carbon sources into distinct entry points within central metabolism (see **Fig. 1**) offer the potential to increase titers, rates, and yields without trade-offs such as nutrient auxotrophy, decreased growth rates, strain stability, and poor energy generation.

**Fig. 1.**
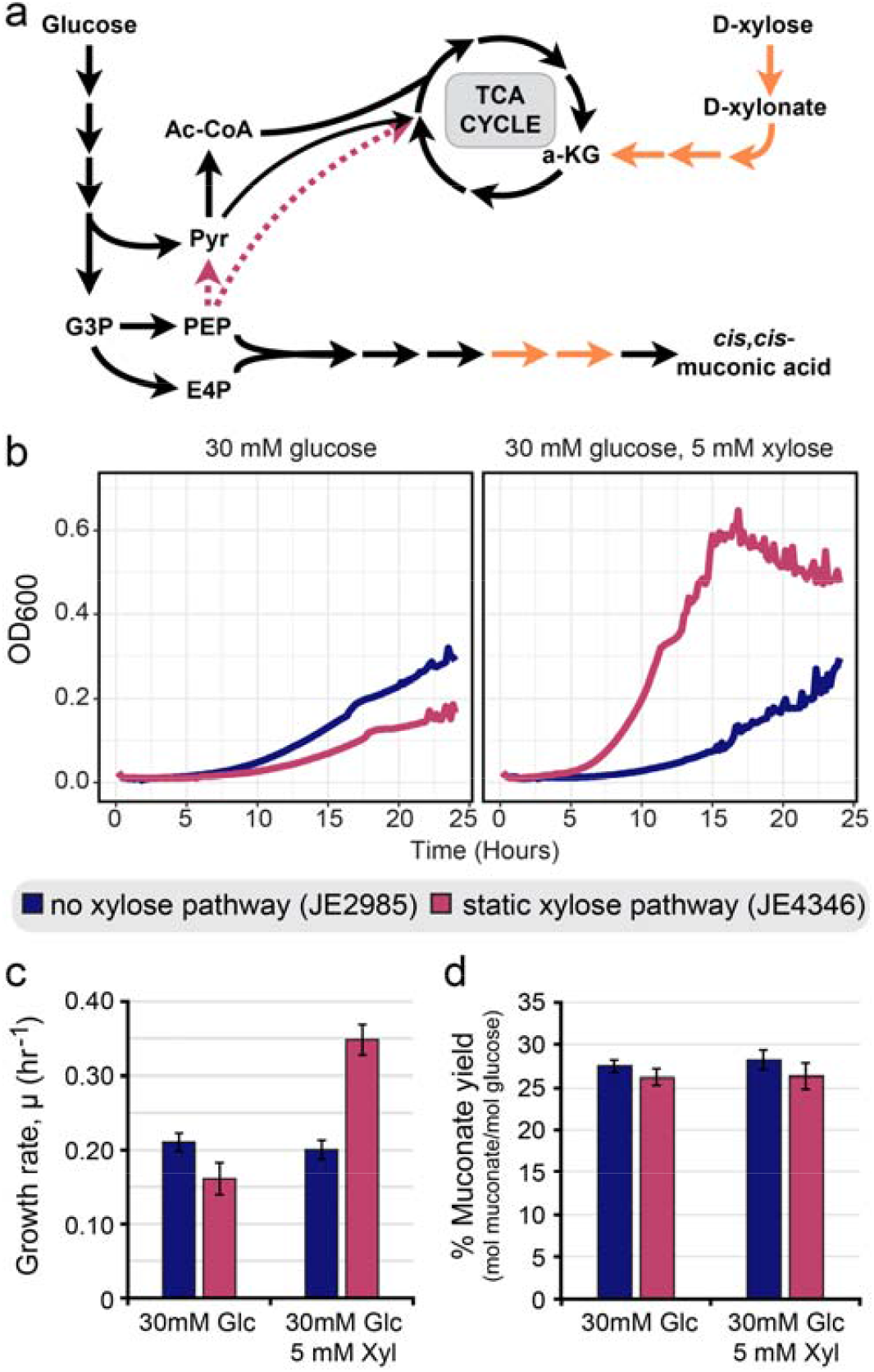
Expression of a heterologous xylose catabolism cassette provides conditional benefits. **(a)** Simplified glucose and xylose catabolic pathways in *Pseudomonas putida* KT2440 with deleted steps indicated by dotted red arrows, heterologous steps indicated by orange arrows, and key metabolites outlined. **(b)** Representative growth curves of sextuplicate cultures for muconic acid production strains in 48-well microtiter plates. Cell density, as measured by OD600, was measured every 10 minutes. **(c)** Growth rates in 48-well microtiter plate cultures for *cis,cis*-muconic acid production strains with and without a static xylose catabolic pathway expression cassette. Strains are grown using glucose or a combination of glucose and xylose as sole carbon sources. **(c)** Molar yield of *cis,cis*-muconic acid per glucose molecule for cultures used to determine growth rates in panel **(b). (b**,**c)** Data are shown as mean values ± standard deviation in six replicates. Abbreviations: G3P (glyceradehyde-3-phosphate), PEP (phosphoenolpyruvate), E4P (erythrose-4-phosphate), 2KG (2-ketoglutarate), Ac-CoA (acetyl-CoA), Pyr (pyruvate), Glc (glucose), Xyl (xylose)

Limitations of current synthetic biology approaches that focus on static expression systems hamper efforts to expand catabolic capabilities beyond a few pathways. These static expression systems are designed for static operating environments, hence they are inefficient in dynamic environments, such as those generated as a consequence of varying substrate consumption kinetics in lignocellulosic bioconversions. The majority of engineered functions, such as inhibitor detoxification (e.g., via catabolism or efflux) and utilization of minor lignocellulosic sugars, are only useful under a subset of environmental conditions. During bioconversion, the catabolic enzymes are no longer needed when the target carbon source is depleted; hence, static expression of these pathways diverts valuable resources away from bioproduct synthesis, reducing carbon efficiency gains afforded by the catabolic repertoire expansion. Despite being masked under many demonstrative conditions (e.g. single pathway expression in a wild-type strain), the metabolic burden of expressing a moderate amount of heterologous pathways can significantly reduce growth rate of otherwise unmodified organisms^10^ and even a single heterologous pathway can substantially reduce performance of an engineered bioproduction strain^11^.

The application of synthetic transcription factor-based biosensors that sense and regulate gene expression in response to specific substrates for dynamic control of engineered catabolic pathways has potential to minimize metabolic burden associated with static gene expression. In recent years, biosensors have been applied with great success to dynamically balance flux through biosynthetic pathways or select for improved strains and enzymes^12-20^. For example, expression of heterologous enzymes in chemical production pathways can be dynamically regulated by feedback inhibition loops that prevent accumulation of undesirable pathway intermediates^14-17^. However, dynamic control strategies have not been directed towards reducing the metabolic burden of expanding the catabolic repertoire of bioproduction strains with heterologous pathway expression.

As a proof-of-concept, in this study we develop a dynamic control system using a novel synthetic xylose biosensor to autonomously regulate expression of a heterologous xylose catabolic pathway in *Pseudomonas putida* – a promising bioproduction platform organism that does not natively catabolize xylose. Previously, *P. putida* has been engineered with static catabolic pathway expression modules that enable use of xylose as a sole carbon source^21-26^. Here we identify and employ a metabolic engineering strategy where synergies arising from engineered glucose and xylose co-utilization improve strain growth and productivity by >65% when compared to utilization of glucose alone. By employing a dynamic control strategy to limit engineered xylose catabolic functions to when xylose is present, we abolish the substantial growth defects that arise from unnecessary pathway expression in the absence of xylose. For this, we develop a xylose biosensor with an array of transcriptional output modalities and optimized ligand binding parameters that enable dynamic control of the metabolic pathway.

## RESULTS

### Constitutive expression of conditionally beneficial catabolic pathways unnecessarily burdens engineered host organisms

Recently, *Pseudomonas putida* KT2440 has been engineered to convert glucose into valuable platform chemicals such as *cis,cis*-muconic acid (ccMA)^27, 28^. High product yields were achieved by redirecting metabolic flux via introduction of genetic modifications that effectively linearize the glucose catabolic pathway, disconnect PEP catabolism from the TCA cycle, and connect the shikimate pathway with truncated aromatic catabolism pathways – resulting in accumulation of valuable aromatic catabolic intermediates (**Fig. 1a**). However, product formation in each of these strains is coupled with growth and glucose catabolism^27^, and the growth rate must be substantially improved to achieve the high volumetric productivity required for industrial processes^28^. We hypothesized that production of TCA cycle intermediates, as well as NAD(P)H production, may be limiting growth in these strains. If correct, then the simultaneous catabolism of glucose with relatively low concentrations of substrates that feed directly into the TCA cycle should improve growth rate. The lignocellulosic pentoses xylose and L-arabinose can be converted to the TCA cycle intermediate 11-ketoglutarate through a linear oxidative pathway called the Weimberg pathway that bypasses traditional glycolysis and produces 2 NAD(P)H molecules per pentose prior to entry into the TCA cycle^29^. Metabolites from the TCA cycle do not readily flow into the pentose phosphate pathway in these strains, so we predict that catabolism of these pentoses with an oxidative pathway will not increase ccMA titers. However, incorporation of these pathways into a ccMA production strain (Fig. 1a) could substantially improve growth rate and productivity in the presence of the corresponding pentose. Finally, given the limited cellular resources available in the engineered ccMA production strains, we also hypothesize that expression of these heterologous pathways under conditions where the pentoses are not present will reduce the fitness of the host.

To test these hypotheses, we integrated a constitutively expressed oxidative xylose catabolism cassette (**Supplementary Fig. S1**) into the genome of the *cis,cis*-muconic acid production strain JE2985, a derivative of CJ442^27^, generating strain JE4346. Specifically, the cassette was designed to constitutively express codon-optimized versions of the xylose:H^+^ symporter (xylE) from *Escherichia coli*^23^, the oxidative xylose catabolism pathway (*xylBCDE*) from *Burkholderia xenovorans*, and 2-ketoglutarate semialdehyde dehydrogenase (*araE*) from *Burkholderia ambifaria* ^29^. We assayed both the growth rate and ccMA production of these strains when cultivated in M9 mineral medium containing glucose with and without supplementary xylose. The growth rate of the parent production strain JE2985 (0.211 hr^-1^) with glucose as a sole carbon source was roughly a 1/4^th^ of wild-type *P. putida* (0.82 hr^-1^), and was not affected by the presence of xylose (**Fig. 1a,b**). As hypothesized, the simultaneous catabolism of glucose and xylose by JE4346 improved growth by >65% over the parent strain (0.348 hr^-1^). However, the metabolic burden of the unnecessary heterologous pathway expression reduced the growth rate by 23% (0.161 hr^-1^) in the absence of xylose. Finally, as predicted, the supplemental xylose had no impact on muconate yield (**Fig. 1c**).

### Development of a xylose biosensor for catabolic pathway control

In nature, expression of metabolic pathways is tuned to provide the ideal amount of enzymatic activity for the prevailing conditions. For example, the soluble components of a regulated catabolic pathway may need high level expression, but in the absence of substrate are unnecessary and near-complete gene silencing may be ideal (**Fig. 2a**). Conversely, substrate transporters often require less expression, and in the absence of substrate some leaky expression may be ideal to facilitate future uptake of the inducing substrate (**Fig. 2a**). Gene expression can be modulated via many mechanisms (*e*.*g*. RBS tuning, controlled protein degradation, promoter modification), and engineering regulated promoters allows tuning of both the leakiness of a promoter as well as the magnitude of induced expression.

**Fig. 2.**
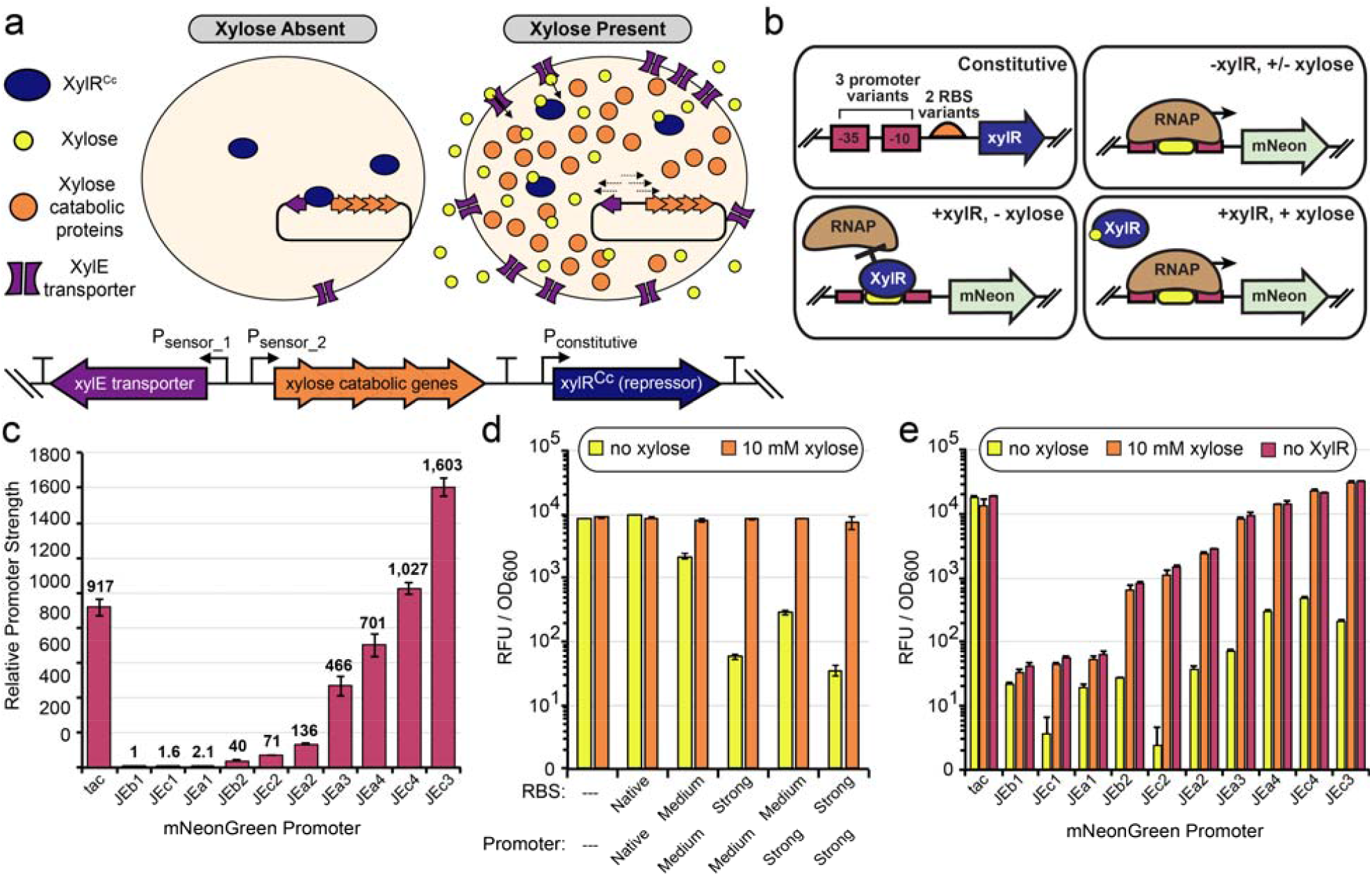
Development of a chromosomally integrated xylose biosensor appropriate for dynamic control of a catabolic pathway. **(a)** Conceptual diagram of a cell utilizing the XylR^Cc^ repressor to employ dynamic control of a xylose catabolic pathway in the presence and absence of xylose. **(b)** Conceptual diagram of XylR^Cc^ expression cassette variants and regulation of fluorescent reporter gene with a xylose-sensitive promoter. **(c)** Graph of relative promoter strengths of the synthetic tac promoter and a small library of native and engineered *Caulobacter crescentus* promoters (JEx) in *P. putida* JE212 which lacks XylR^Cc^. **(d)** Graph of mNeonGreen production by strains containing both XylR^Cc^ expression variants and an integrated P_JEa3_:mNeonGreen cassette during growth in mineral media containing glucose ± xylose. The ribosomal binding site (RBS) and transcriptional promoter variants for XylR expression are indicated on the x-axis. **(e)** Graph of mNeonGreen production in JE212 (red) or XylR^Cc^ -containing JE2926-variants (orange and yellow) with integrated mNeonGreen promoter fusion variants in the presence (orange) or absence of xylose (yellow and red) of xylose. **(c-e)** Background RFU/OD600 fluorescence from a strain lacking a mNeonGreen cassette is subtracted to generate relative promoter strength values. Data are presented as the mean values ± standard deviation in three replicates.

Because the heterologous pathway was beneficial in the presence of xylose but detrimental in the absence, we aimed to mimic nature and use dynamic control to autonomously regulate expression of the catabolic pathway to limit expression when it is beneficial. To accomplish this, we first designed a chromosomally integrated xylose biosensor that would enable dynamic control of heterologous xylose catabolic pathway expression. When selecting candidate xylose biosensors, we avoided transcriptional activators^30^ due to the complexity of downstream promoter engineering and repressors that interact with known anti-inducers such as glucose^31^. Furthermore, pathway optimization requires promoters with diverse transcriptional outputs, and the xylose-sensitive transcriptional repressor from *Caulobacter crescentus* NA1000 (XylR^Cc^) regulates several promoters^32^, so it was chosen as the basis for the xylose biosensor.

We generated and assayed performance of a small library of σ^70^ promoters based upon three native xylose-regulated C. crescentus promoters, two from *C. crescentus* NA1000 (P_*JEa1*_ and P_*JEb1*_) and one from C. crescentus K31 (P_*JEc1*_). Promoters were assayed first in a strain lacking XylR^Cc^ to determine the maximal expression level in the fully unrepressed state using mNeonGreen (**Fig. 2b,c**). This was performed in *P. putida* strain JE212, a KT2440 derivative containing both a genome integrated copy of the Bxb1 integrase and deletion of *gcd* to prevent conversion of xylose to xylonate in the periplasm^23^. Based on prior analysis of how -35 and -10 sequences correlate with σ^70^ promoter strength in *P. putida*^33^, and the fact that UP-elements that can compensate for -10 and -35 sequences^33, 34^ are poorly conserved between bacterial species, we predicted that the *C. crescentus* promoters may provide weak expression in *P. putida*. Accordingly, we observed that the tac promoter, which is a strong promoter in *P. putida*^33^, enabled mNeonGreen production levels 446-to 917-fold higher than the *C. crescentus* promoters (**Supplementary Table S1, Fig. 2c**). We therefore modified the -35 and -10 sequences for each native promoter to expand the range of promoter outputs, resulting in promoters P_*JEa2*_, P_*JEb2*_, P_*JEc2*_, P_*JEa3*_, P_*JEb3*_, and P_JEc3_. These promoters provide a range of unrepressed expression levels covering 3 orders of magnitude and serve as biosensor output modules that can be used to tune expression of the xylose catabolic pathway components.

We next assessed the ability XylR^Cc^ to regulate each of the promoters from the library. A codon-optimized version of *xylR*^Cc^ under control of its native upstream regulatory sequence was integrated into the chromosome of JE1603 – a derivative of JE212 that includes a chromosomally integrated xylose H^+^/symporter. However, the native expression of XylR^Cc^ was insufficient to repress expression of mNeonGreen by the P_JEa3_ promoter in the absence of xylose (**Fig. 2d**). We hypothesized that insufficient XylR^Cc^ expression explained the lack of repression, and constructed four additional XylR^Cc^ cassettes (**Fig. 2b**) with four promoter and ribosomal binding site (RBS) combinations predicted by our previous work^33^ and the Salis Lab RBS calculator^35^ to provide a 56-fold range of XylR^Cc^ expression (**Table 1**). Each of the four XylR^Cc^ expression cassettes supported repression of P_JEa3_, and the RBS sequence was critical for robust repression. In comparison with the *medium* RBS, utilizing a strong RBS for XylR^Cc^ increased the degree of repression by 38.6-fold (medium promoter) and 10-fold (strong promoter). Increasing the transcription of *xylR*^Cc^ had little impact on repression when utilizing the *strong* RBS, suggesting that expression of XylR^Cc^ is approaching saturation. Accordingly, we selected the *medium* promoter, *strong* RBS *xylR*^Cc^ cassette in JE2926 for subsequent experiments because its lower XylR^Cc^ expression should minimize the metabolic burden of repressor expression.

**Table 1.**
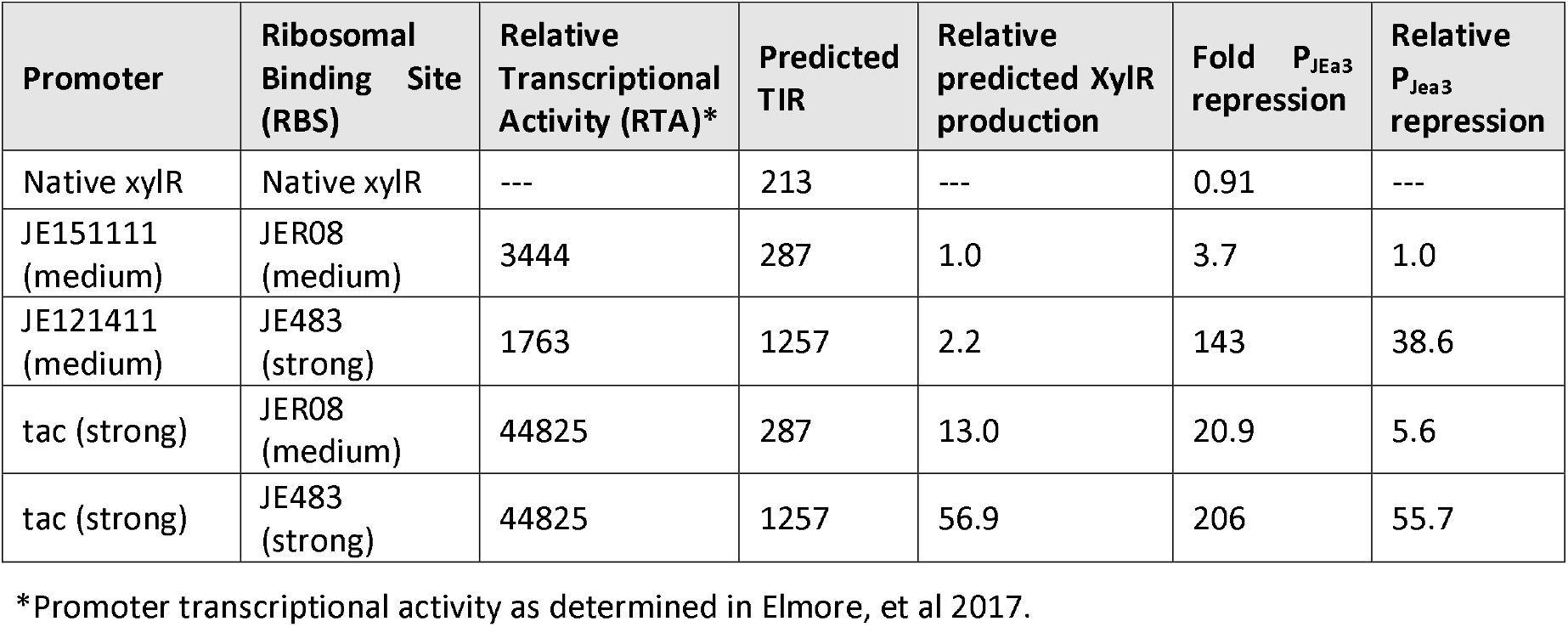
Performance of XylR^Cc^ expression cassettes.

We next evaluated whether the remaining members of the promoter library are regulated in the presence of xylose in JE2926 (**Fig. 2e**). In the presence of 10 mM xylose (**Fig. 2e, orange bars**) the performance of each promoter was indistinguishable from its performance in the absence of XylR^Cc^ (**Fig. 2d, red bars**), indicating complete derepression by xylose. However, in the absence of xylose, each promoter displayed varying degrees of repression (**Fig. 2d, yellow bars**). The largest fold-repression was observed with the moderately strong promoters P_JEa3_ (118-fold) and P_JEc2_ (457-fold). Stronger variants of these promoters are less well repressed, which may be a result of increased competition with the RNA polymerase (RNAP) for promoter occupancy as seen when comparing *tac* and *lac* promoters in *E. coli*^36^. The degree of repression measured for the weaker promoters is low and is likely underestimated as a consequence of *P. putida* autofluorescence impacting the mNeonGreen limit of detection. Taken together, the optimized xylose biosensor coupled with a wide array of output modules should serve well for dynamic control of xylose catabolic functions.

### Employing dynamic control for regulation of a heterologous xylose catabolic pathway

We next sought to determine if the optimized xylose biosensor and collection of regulated promoters would enable dynamic control of the xylose isomerase catabolic pathway that we previously developed for *P. putida* KT2440 ^23^ (**Fig. 3a**). The original cassette contained two operons under control of static expression systems. The optimized XylR^Cc^ cassette from JE2926 was inserted downstream of the catabolic gene operon, and the static promoters controlling each operon (P_trans_ and P_tac_) were systematically replaced with xylose-sensitive promoters. Each pathway variant was incorporated into the *aldB-I* locus of JE212, and the resulting strains assayed for growth with xylose. As expected, the growth rate with the unmodified expression system (0.163 hr^-1^) was similar to the previously reported growth rate (0.150 hr^-1^), which suggests that chromosomal location and incorporation of XylR^Cc^ had little impact on pathway performance.

**Fig. 3.**
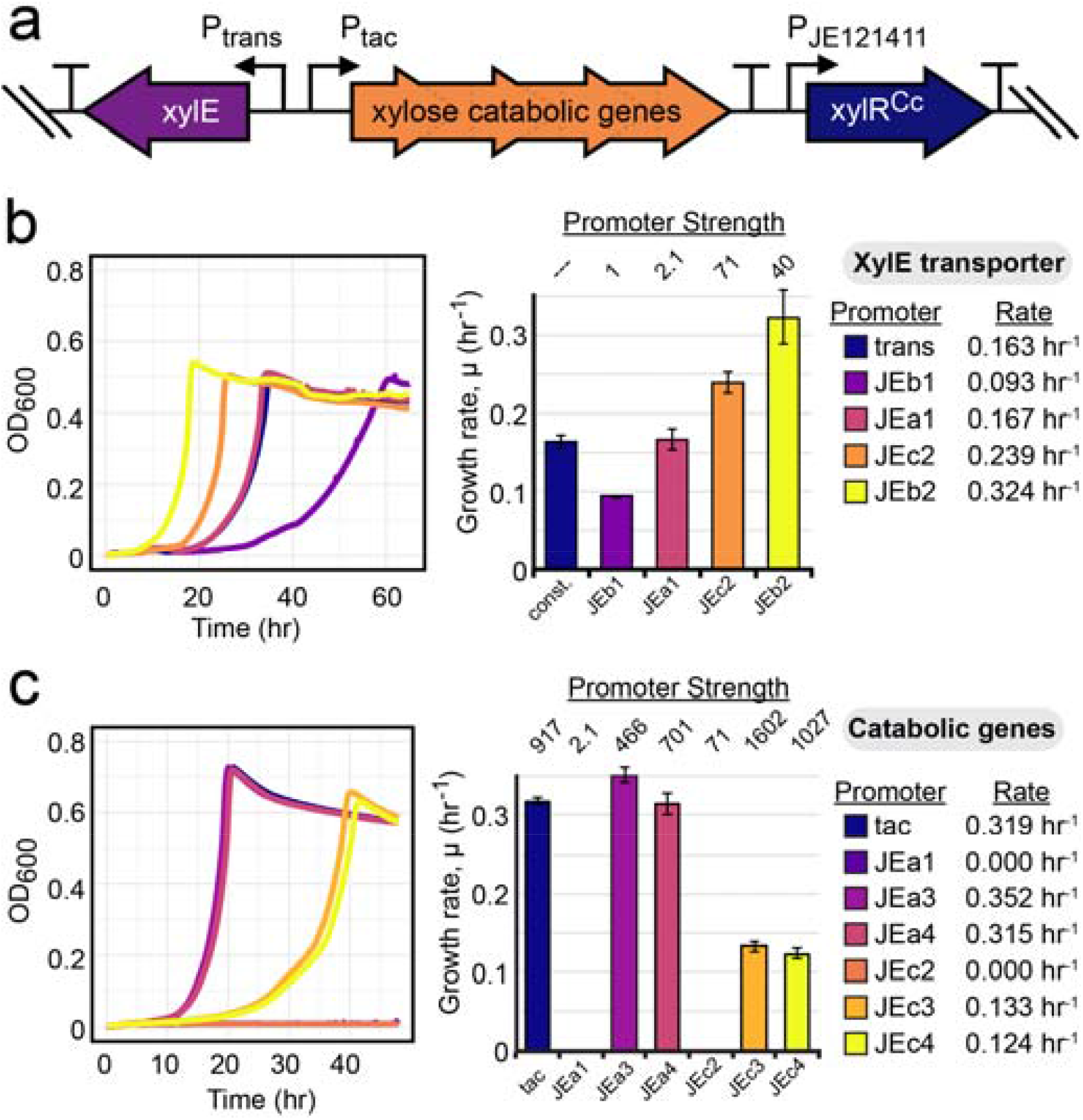
Optimizing dynamic control of a heterologous xylose isomerase pathway in *Pseudomonas putida*. **(a)** Diagram of gene arrangement in the synthetic xylose catabolism cassette containing XylR^Cc^. Promoters and rho-independent terminators are indicated using black arrows and T’s respectively. **(b**,**c)** Representative growth curves and growth rates from cultures with xylose as sole carbon source in 48-well microtiter plate cultivations. Cell density, as measured by OD600, was measured every 10 minutes. Strains in which static promoters for **(b)** XylE transporter **(c)** catabolic pathway genes have been replaced with xylose-regulated promoters are indicated by color. Bar graph growth rate data are represented as the mean values ± standard deviation in three replicates.

Replacing the static promoters with dynamically regulated promoters substantially improved to growth rate on xylose. Tuning transporter expression over a 71-fold range of expression (**Fig. 3b**) doubled the growth rate (0.163 hr^-1^ vs. 0.324 hr^-1^), and a further 10% increase in growth rate was achieved when catabolic operon expression was reduced by ∼50% (**Fig. 3c** – P_tac_ vs P_JEa3_). In both cases using the strongest promoter provided suboptimal performance – particularly in the case of the catabolic pathway, where a weaker promoter (P_JEb3_) enabled a growth rate 2.8-fold higher than the strongest promoter (P_JEc3_). We also confirmed that XylR^Cc^ was functional in each strain by assessing xylose-sensitivity of a P_JEa3_:*mNeonGreen* reporter that was chromosomally integrated into each strain (**Supplementary Fig. S2**).

### Engineering ligand sensitivity and specificity of XylR^Cc^ for lignocellulosic pentose sugars

In natural environments, carbon sources are often found in very low concentrations that are significantly lower than those used in laboratory or industrial scale processes. Accordingly, the associated transcription factors for such carbon sources frequently respond to similarly low concentrations (e.g., high nM to low μM)^37-40^. Furthermore, transcription factors can often interact with molecules that are structurally similar to the target metabolite, albeit at lower sensitivity. Thus it is not surprising that the XylR^Cc^ transcription factor responds to concentrations as low as 2 μM xylose in *C. crescentus*^37^. *C. crescentus XylR*^Cc^ was also demonstrated to weakly interact with another major lignocellulosic pentose, L-arabinose^32^. To examine the sensitivity and specificity of XylR^Cc^ towards xylose in our system, we tested the expression of a P_JEa2_:mNeonGreen cassette in *P. putida*. Complete derepression was attained with a xylose concentration of 10 μM xylose (**Fig. 4a**). We also assayed activation mNeonGreen in the presence of 1 μM to 10 mM L-arabinose (**Fig. 4a**). In accordance with the previous report, XylR^Cc^ was also derepressed by L-arabinose, but required >100-fold more L-arabinose (1 mM) to achieve maximal mNeonGreen expression. While 1 mM (150 mg/L) L-arabinose concentrations may be uncommon nature, such concentrations will be present in many lignocellulosic hydrolysates^7, 41-43^.

**Fig. 4.**
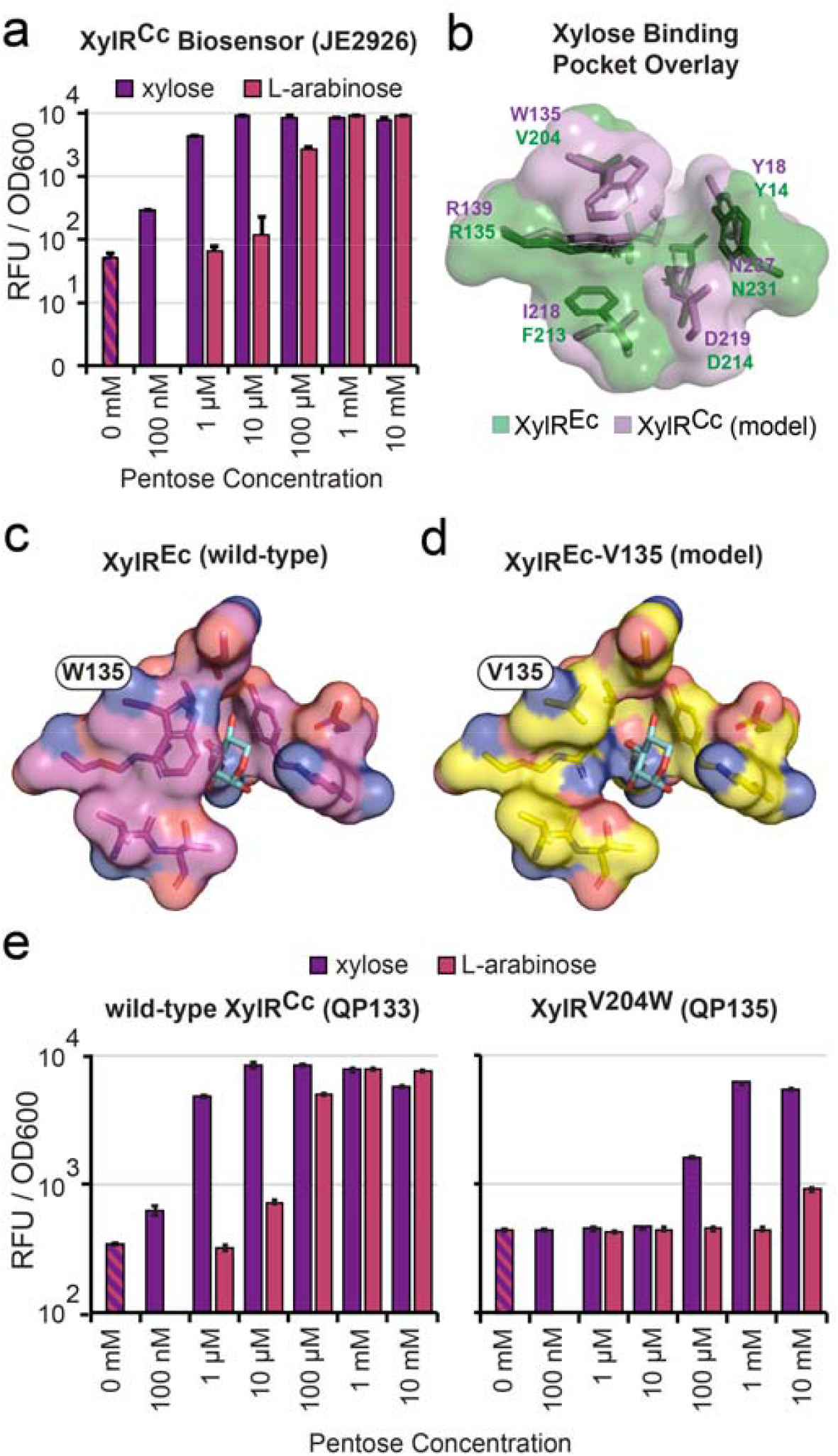
Engineered steric hindrances in the substrate binding pocket reduces derepression of XylR^Cc^ by L-arabinose. **(a**,**e**,**f)** Graph of mNeonGreen production by xylose biosensor strains when exposed to no inducer (red and purple striped), xylose (purple), or L-arabinose (red) during growth with glucose as sole carbon source in microtiter plate cultivations. **(b)** XylR^Cc^ structure modeled upon PDB ID: 4FE4 overlayed onto structure 4FE4 of *Escherichia coli* XylR (XylR^Ec^). Wild-type **(c)** and modeled W135V mutant **(d)** structures of XylR^Ec^ with bound xylose ligand (PDB ID:4FE7). Images **(b-d)** and models structures **(b**,**d)** were generated using PyMol 2.2.3. **(a**,**e)** Data are presented as the mean values ± standard deviations in three replicates.

Given the demonstrated metabolic burden of unnecessary catabolic pathway expression (**Fig. 1**), we sought to engineer a XylR^Cc^ variant with reduced sensitivity towards L-arabinose to prevent inadvertent activation of the xylose pathway expression by L-arabinose. We used modeled structural information to guide rational engineering of XylR^Cc^ ligand binding. Solved structures are not available for XylR^Cc^, so structures of the xylose-sensitive transcriptional activator from *E. coli* (XylR^Ec^) were used with and without bound ligand as references for modeling a XylR^Cc^ structure. Unlike XylR^Cc^, Ni and colleagues reported that the XylR^Ec^ activator does not interact with L-arabinose^40^. Furthermore, XylR^Ec^ requires >200-fold more (>2 mM) xylose than XylR^Cc^ for maximal activation of reporter gene expression^44^. The modeled structure of XylR^Cc^ contains a central region where several of the residues that interact with xylose in XylR^Ec^ are positioned similarly to the modeled XylR^Cc^ structure (**Fig. 4b, Supplementary Fig. S3**). While W135 does not directly interact with xylose (**Fig. 4c**), it helps define the binding pocket, and a steric clash between W135 and L-arabinose in XylR^Ec^ is thought to impact binding^40^. However, a much smaller valine residue (V204) occupies this position in XylR^Cc^ and may result in a more accessible ligand binding pocket. Indeed, when the tryptophan is replaced with a valine in XylR^Ec^ the binding pocket is more exposed (**Fig. 4d**). Accordingly, we hypothesized that replacing V204 in XylR^Cc^ with tryptophan would introduce a steric hindrance in the binding pocket that both reduces sensitivity for pentoses and further increases its specificity for xylose.

To compare wild-type *xylR* (Xyl^Cc^) with the mutant allele (XylR^V204W^), we integrated synthetic modules containing operons for either wild type or V204W xylR with P_JEa3_:*mNeonGreen* (**Supplementary Fig. S4**) into JE4681, which contains codon-optimized *E. coli* L-arabinose and D-xylose symporters that have been previously validated in *P. putida*^23^. The resulting strains QP133 and QP135, expressing XylR^Cc^ and XylR^V204W^, respectively, demonstrated substantially different responses to xylose and L-arabinose. Aside from a higher basal level of expression – potentially a result of XylR^Cc^ repression via run-on transcription – the performance with wild-type XylR^Cc^ (**Fig. 4e**) was essentially identical with our previous results (**Fig. 4a**). However, activation of gene expression with XylR^V204W^ required a two- and three-order of magnitude increase in ligand concentration for xylose and L-arabinose, respectively. Derepression of XylR^V204W^ by L-arabinose was not observed below 10 mM (1.5 g/L), and thus the use of XylR^V204W^ should limit undesired activation of xylose pathway expression in lignocellulosic hydrolysates depleted of xylose.

We next assayed XylR^V204W^ for dynamic control of xylose catabolism. For this, the *xylR* mutation was introduced into JE3238, which contains the best performing xylose isomerase pathway cassette, generating strain JE4280. Versions of the two strains containing a chromosomally-integrated P_JEa3_:mNeonGreen cassette were assayed for growth with 10 mM glucose, 10 mM xylose, or 5 mM of each carbon source. The growth rate and lag phase were indistinguishable when xylose or glucose were the sole carbon source (**Supplementary Fig. S5, Supplementary Fig. S6**). As expected, mNeonGreen was repressed when glucose was the sole carbon source (**Supplementary Fig. S6a**,**c**). While both strains perform identically during the early-to-mid cultivation with mixed carbon sources, the strain utilizing the lower sensitivity XylR^V204W^ repression displays reduced mNeonGreen expression and lower growth rate near the end of the cultivation (**Supplementary Fig. S6b**,**d**). A possible explanation is that when xylose nears depletion in the medium the intracellular concentrations of xylose are insufficient for full derepression of XylR^V204W^, but not wild-type XylR^Cc^, regulons. Because glucose is consumed faster than xylose by engineered *P. putida*^23^, xylose is likely the sole carbon source near the end of cultivation in media with mixed carbon sources, and reduced catabolic pathway expression would thus lead to a reduced growth rate. During industrial cultivations xylose concentrations will generally be well above this threshold, higher density cultures will more rapidly consume residual xylose, and rapid repression of the xylose regulon upon depletion of the carbon source would likely be beneficial.

### Dynamic control abolishes the burden of a heterologous xylose catabolic pathway in a *cis,cis*- muconic acid production strain

We finally sought to determine whether dynamic control of xylose catabolism would eliminate the growth defect (**Fig. 1b**) associated with catabolic protein expression under conditions where expression is unnecessary. We used three synthetic modules containing the previously described oxidative xylose catabolic cassette (**Supplementary Fig. S1**) that differ only by the presence or absence of a *xylR* operon to enable dynamic regulation of the transporter and catabolic pathway. These modules were first integrated into the non-bioproduction strain JE212 to assay performance of the pathways. As observed with the xylose isomerase pathway (**Supplementary Fig. S5**), growth rates when using xylose as a sole carbon source were very similar (**Supplementary Fig. S7**), regardless of whether expression of the xylose catabolism cassette was static (0.181 hr^-1^) or dynamically controlled by XylR^Cc^(0.171 hr^-1^) and XylR^V204W^ (0.176 hr^-1^).

Given the similar performance of the dynamically controlled and static xylose catabolic cassettes pathways in JE212, we hypothesized that a strain with the dynamically controlled cassette would provide the benefits of the pathway when xylose is present, without the detriment of the pathway in its absence. We integrated the dynamically controlled xylose catabolic cassette into the *cis,cis*-muconate production strain JE2985, generating strain JE4349. It is important to note that JE4349, aside from the presence of the *xylR* operon, is isogenic with JE4346, the JE2985-derivative containing a constitutive oxidative xylose catabolic pathway (**Fig. 1**). We tested performance of all three strains in minimal media containing 30 mM glucose with or without 5 mM supplementary xylose (**Fig. 5a**). As hypothesized, when the medium contains xylose, the growth rates of each xylose-catabolizing strain >60% faster than the parent strain (Fig. 5b). However, dynamic control of the xylose catabolism cassette expression enabled ∼30% faster growth with glucose as the sole carbon source – effectively abolishing the fitness defect imposed by static expression of the catabolic genes. Finally, under both growth conditions, muconic acid was similar between each of the three strains (**Fig. 5b**). Taken together, this demonstrates that unnecessary expression of heterologous catabolic pathways incurs a fitness cost and highlights the utility of dynamic pathway regulation to abolish fitness defects while maintaining the benefits of the catabolic pathway.

**Fig. 5.**
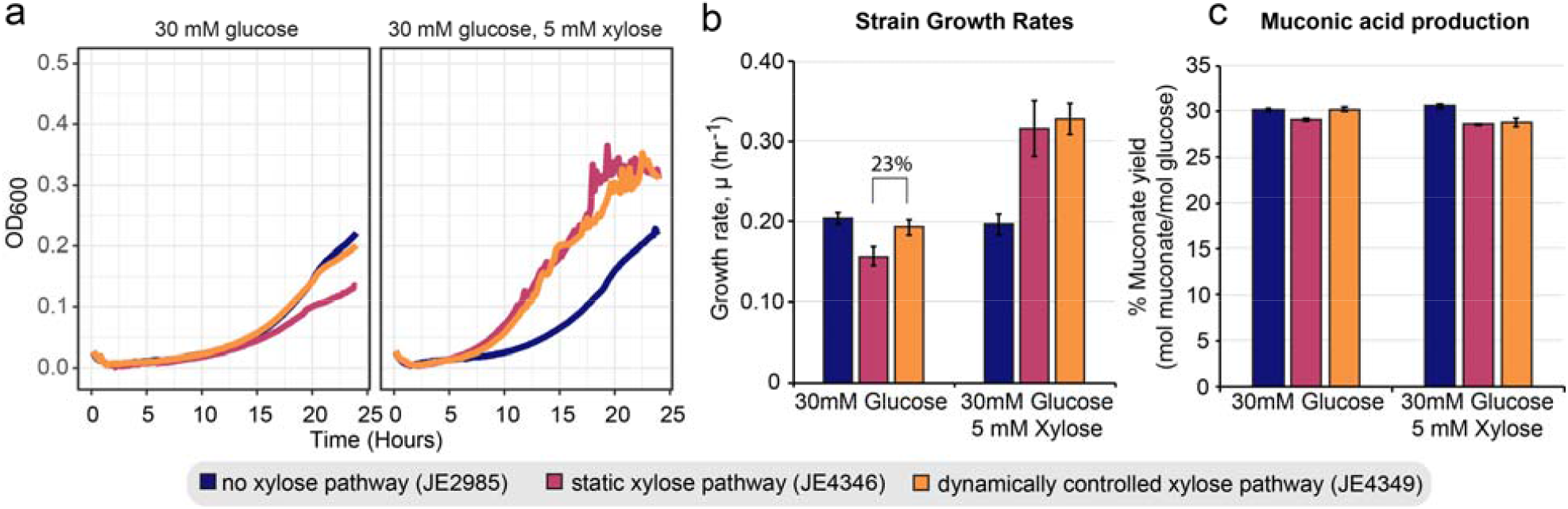
Dynamic control of heterologous catabolic pathways abolishes the fitness defects that arise from unnecessary static pathway expression. **(a)** Representative growth curves of sextuplicate cultures for muconic acid production strains in 48-well microtiter plates. Cell density, as measured by OD600, was measured every 10 minutes. **(b)** Growth rates in 48-well microtiter plate cultures for *cis,cis*-muconic acid production strains with and without a static xylose catabolic pathway expression cassette. Strains are grown using glucose alone or a combination of glucose and xylose as sole carbon sources. **(c)** Molar yield of *cis,cis*-muconic acid per glucose molecule for cultures used to determine growth rates in panel **(b). (b**,**c)** Data are represented as mean values ± standard deviation in six replicates. Source data underlying Fig. 5 are provided as a Source Data file.

## DISCUSSION

This work describes the development of an approach for establishing dynamic control of heterologous catabolic pathways and provides a template for future efforts to engineer chromosomally integrated dynamic control systems directly in emerging microbial hosts. We define and employ criteria for selecting transcription factors that are appropriate for dynamic control and utilize synthetic biology approaches to optimize expression of a single copy, heterologous transcription factor. Additionally, we utilized rational design to generate a small collection of xylose-sensitive promoters with a range of outputs ranging over three orders of magnitude and modify the ligand binding parameters of the associated transcription factor to change the dynamic range.

As the replicating plasmids frequently used to host biosensors are unsuitable for deployment in industrial bioreactors, plant rhizospheres, or human microbiomes the rapid optimization of transcription factor and output promoter performance in host chromosomes will be critical for developing dynamic control systems. We envision future dynamic control system design will combine methodologies that were first utilized for biosensor development in *E. coli*^20^ (*e*.*g*. targeted promoter and transcription factor mutagenesis, high-throughput promoter screening, fluorescence-assisted cell sorting) with modern host-agnostic genome engineering tools (e.g. SAGE, CRAGE)^45, 46^ to enable rapid plasmid-free biosensor development directly in diverse microbial host chromosomes. Combining these approaches to generate and test large combinatorial libraries of parts will enable rapid biosensor development with minimal *a priori* knowledge.

Robust co-utilization of multiple carbon sources will be critical for carbon efficient bioconversion of complex feedstocks and will enable novel metabolic engineering strategies that are impossible when considering bioconversion of single carbon sources. Here, we introduced an oxidative xylose catabolism pathway into *Pseudomonas putida* that has been heavily engineered for high yield bioconversion of glucose to ccMA^27^. Conversion of a relatively small amount of xylose drastically improved the growth rate of the strain with minimal change to yield of *cis,cis*-muconic acid from glucose. As a consequence of its rearranged central metabolism, the oxidative xylose catabolism pathway is unable to support growth on xylose as a sole carbon source in the bioproduction strain. Thus, synergy between glucose and xylose catabolic pathways, rather than rapid catabolism of xylose as a sole carbon source, is responsible for the improved performance. A similar strategy could be employed using oxidative pathways for L-arabinose or galacturonic acid^29^. This may even be preferable, as sacrificing these lower abundance sugars for energy and biomass production would free the more highly abundant xylose for bioconversion into products. In this scenario, dynamic control would be even more critical, as these sugars are likely to depleted earlier in a bioprocess and their presence is less universal among lignocellulosic feedstocks. Successful demonstrations of engineered carbon source co-utilization^10, 23, 47-51^ in recent years have paved the way for deploying novel metabolic engineering strategies centered around co-utilization of multiple substrates in complex feedstocks such as lignocellulosic hydrolysates^52^. Ultimately, strategies such as dynamic pathway control that reduce the metabolic burden of heterologous catabolic pathway expression will be essential for extensive expansion of an organism’s catabolic repertoire.

## ACKNOWLEDGEMENTS

This work was authored in part by Oak Ridge National Laboratory, which is managed by UT-Battelle, LLC, for the U.S. Department of Energy under contract DE-AC05-00OR22725. This work was also partially authored under Triad National Security, LLC (“Triad”) Contract No. 89233218CNA000001 with the U.S. Department of Energy and Battelle, the manager and operator of the Pacific Northwest National Laboratory for the U.S. Department of Energy under Contract No DE-AC05-76RLO1830. This research used computational resources provided by the Los Alamos National Laboratory Institutional Computing Program to R.K.J., which is supported by the U.S. Department of Energy National Nuclear Security Administration under Contract No. 89233218CNA000001, Research was in part sponsored by the Laboratory Directed Research and Development Program of Oak Ridge National Laboratory, Project ID 7866. We also acknowledge funding from the U.S. Department of Energy Office of Energy Efficiency and Renewable Energy the Bioenergy Technologies Office via the Agile BioFoundry project.

## MATERIALS AND METHODS

### General culture conditions & media

The strains and plasmids used in this study are listed in Supplemental Table S2. Routine cultivation of *Escherichia coli* for plasmid construction and maintenance was performed at 37 °C using LB (Miller) medium supplemented with 50 μg/mL kanamycin sulfate and 15 g/L agar (for solid medium). All *Pseudomonas putida* assay cultures were incubated at 30 °C, and 250 rpm with 10 mM orbital for cultures performed in shake flasks (strain maintenance, competent cell preparations, and starter cultures) and 548 rpm with a 2 mM orbital (fast shaking setting) for cultures performed in plate readers from BioTek - Neo2SM plate reader (fluorescent reporter assays) or Epoch2 plate reader (growth rate determination, ccMM production assays). BCDL LB (Miller) was used for routine *Pseudomonas putida* strain maintenance, competent cell preparations, and starter cultures.

MME medium (containing 9.1 mM K_2_HPO_4_, 20 mM MOPS, 4.3 mM NaCl, 9.3 mM NH_4_Cl, 0.41 mM MgSO_4_, 68 μM CaCl_2_, 1x MME trace minerals, pH adjusted to 7.0 with KOH) supplemented with carbon sources as indicated in the test was utilized for biosensor development and pathway optimization assays. Modified M9 medium (47.8 mM Na_2_HPO_4_, 22 mM KH_2_PO_4_, 18.7 mM NH_4_Cl, 8.6 mM NaCl, 1 mM MgCl_2_, 0.1 mM CaCl2, 18 μM FeSO_4_, 1x MME trace minerals, pH adjusted to 7 with KOH) containing 30 mM glucose with or without 5 mM xylose was used for assaying growth rates and product yields from ccMM production strains. 1000x MME trace mineral stock solution contains per liter, 1 mL concentrated HCl, 0.5 g Na_4_EDTA, 2 g FeCl_3_, 0.05 g each H_3_BO_3_, ZnCl_2_, CuCl_2_·2H_2_O, MnCl_2_·4H_2_O, (NH_4_)_2_MoO_4_, CoCl_2_·6H_2_O, NiCl_2_·6H_2_O.

### Plasmid & Pseudomonas strain construction

Phusion→ HF Polymerase (Thermo Scientific) and primers synthesized by Eurofins Genomics were used in all PCR amplifications for plasmid construction. OneTaq® (New England Biolabs - NEB) was used for colony PCR. Plasmids were constructed by Gibson Assembly using NEBuilder® HiFi DNA Assembly Master Mix (NEB) or ligation using T4 DNA ligase (NEB). Plasmids were transformed into either competent NEB 5-alpha F’I^q^ (NEB) or Epi400 (Lucigen). Standard chemically competent *Escherichia coli* transformation protocols were used to construct plasmid host strains. Transformants were selected on LB (Miller) agar plates containing 50 μg/Ml kanamycin sulfate for selection and incubated at 37 °C. Template DNA was either synthesized (IDT or Eurofins) or isolated from *E. coli* or *P. putida* KT2440 using Zymo Quick gDNA miniprep kit (Zymo Research). Zymoclean Gel DNA recovery kit (Zymo Research) was used for all DNA gel purifications. Plasmid DNA was purified from *E. coli* using GeneJet plasmid miniprep kit (ThermoScientific) or ZymoPURE plasmid midiprep kit (Zymo Research). Sequences of all plasmids were confirmed using Sanger sequencing performed by Eurofins Genomics. Plasmids used in this work are listed in Supplemental Table S2, and details regarding plasmid construction and sequences are below.

*P. putida* JE90, a derivative of *P. putida* KT2440 where the restriction endonuclease *hsdR* has been replaced with the Bxb1-phage integrase and respective *attB* sequence^33^ or *P. putida* CJ442, a muconic acid production strain^27^, were used as parents for all *P. putida* strains used in this study (Supplemental Table S2). All genome modifications were performed using either the homologous recombination-based pK18mobsacB kanamycin resistance/sucrose sensitivity selection/counter-selection system^27^ as described in detail previously^23^ or with the Bxb1-phage integrase system^33^ with minor modifications to competent cell preparation procedures as described in detail previously^23^. Primers used for screening *P. putida* strains for *aldB-I* or *ampC* replacement can be found in Supplemental Table S3. Integration of pJE990-derivatives using the phage integrase system was confirmed by colony PCR using oligos oJE66 & oJE536. Annotated sequences of all plasmids used for strain construction can be found in “Supplemental File P1”.

### Growth rate analysis

For non-ccMM production strains: 5 mL LB medium was inoculated from glycerol stocks and incubated overnight at 30°C, 250 rpm for precultures cultures. Starter cultures were prepared by inoculating 5 mL MME supplemented with 10 mM glucose from LB precultures (1% inoculum), and incubated at 30°C, 250 rpm until stationary phase was reached (typically overnight) to synchronize cultures and normalize inoculum. Growth assays were performed with 600 uL MME supplemented with appropriate carbon sources per well in clear 48-well microtiter plates with an optically clear lid (Greiner Bio-One). Starter culture OD600 values were spot-checked to ensure similar starting inoculum. Assay plates were incubated at 30 °C with fast shaking setting in a BioTek plate reader (Bio-Tek), with OD_600_ readings taken every 10 minutes. Exponential growth rates were determined using the CurveFitter software (http://www.evolvedmicrobe.com/CurveFitter/) with data points in mid-log phase where OD_600_ is between 0.04-0.2 (not equivalent to 1 cm path length). All growth rates were calculated from 3 replicates. Growth curves displayed in figures are representative curves derived from one of the replicates.

For ccMM production strains: 50 mL LB medium was inoculated from glycerol stocks, and incubated overnight at 30°C, 250 rpm for starter cultures. Cultures were washed 3x with 25 mL modified M9 medium lacking a carbon source to remove residual LB. Assays were performed with 600 mL modified M9 medium supplemented with appropriate carbon sources per well in clear 48-well microtiter plates with an optically clear lid (Greiner Bio-One). Each culture was inoculated to an initial OD_600_ of 0.1 (as determined using a 1 cm pathlength cuvette). Assay plates were incubated at 30 °C with fast shaking setting in a BioTek plate reader (Bio-Tek), with OD_600_ readings taken every 10 minutes. Exponential growth rates were determined using the CurveFitter software (http://www.evolvedmicrobe.com/CurveFitter/) with data points in mid-log phase where OD_600_ as measured by the plate reader is between 0.04-0.15 (not equivalent to 1 cm path length). All growth rates were calculated from 3 or 4 replicates. Growth curves displayed in figures are representative curves derived from one of the replicates.

## Fluorescent reporter assays

Precultures were generated as described above for non-ccMM production strains. Coupled growth and fluorescence assays were performed with a Neo2SM (Bio-Tek) plate reader using 200 μL/well of MME containing appropriate carbon sources in black-walled, μClear® flat-bottom, 96-well plates (Greiner Bio-One) with an optically clear lid. Plate cultures were inoculated with 0.5% inoculum from starter cultures, and incubated overnight at 30 °C, fast shaking setting with OD_600_ and fluorescence (F_510,530_) measured every 10 minutes. Reporter expression per cell was estimated by dividing relative fluorescence units (RFU) by OD_600_ (as a proxy for cell number) for each time point and averaging those values for time points occurring during mid-log growth phase where OD_600_ as measured by the plate reader is between 0.1-0.2. Background absorbance and fluorescence readings from wells containing a strain containing a promoterless fluorescent reporter were averaged and subtracted from sample readings prior to analysis. Standard deviations are two-sided.

### Analytical techniques

HPLC analysis for glucose, xylose, and ccMM detection was performed by injecting 20 μL of 0.2 μm filtered culture supernatant from end point growth rate analysis cultures onto a Waters 1515 series system equipped with a Rezex RFQ-Fast Acid H+ (8%) column (Phenomenex) and a Micro-Guard Cation H^+^ cartridge (Bio-Rad). Samples were run with column at 60 °C using a mobile phase of 0.01 N sulfuric acid at a flow rate of 0.6 mL/min, with a refractive index detector and UV/Vis detector measuring A_230_ & A_280_ for analyte detection. Analytes were identified and quantified by comparing retention times and spectra with pure standards. The effect of evaporation in microtiter plate wells (and therefor sample concentration) on measurements was compensated for by correcting concentrations using intensity of a spectra generated by a salt in the medium and comparing HPLC results of each sample versus the original medium.

Standard deviations are two-sided.

## Statement on measurements

For all data points in this manuscript, measurements were taken from distinct samples.

