## Supplementary Materials for "Dynamic control systems that mimic natural regulation of catabolic pathways enable rapid production of lignocellulose-derived bioproducts"

**Dynamic control systems that mimic nature enable carbon efficient production of lignocellulose-derived bioproducts**

*Elmore, et al.*

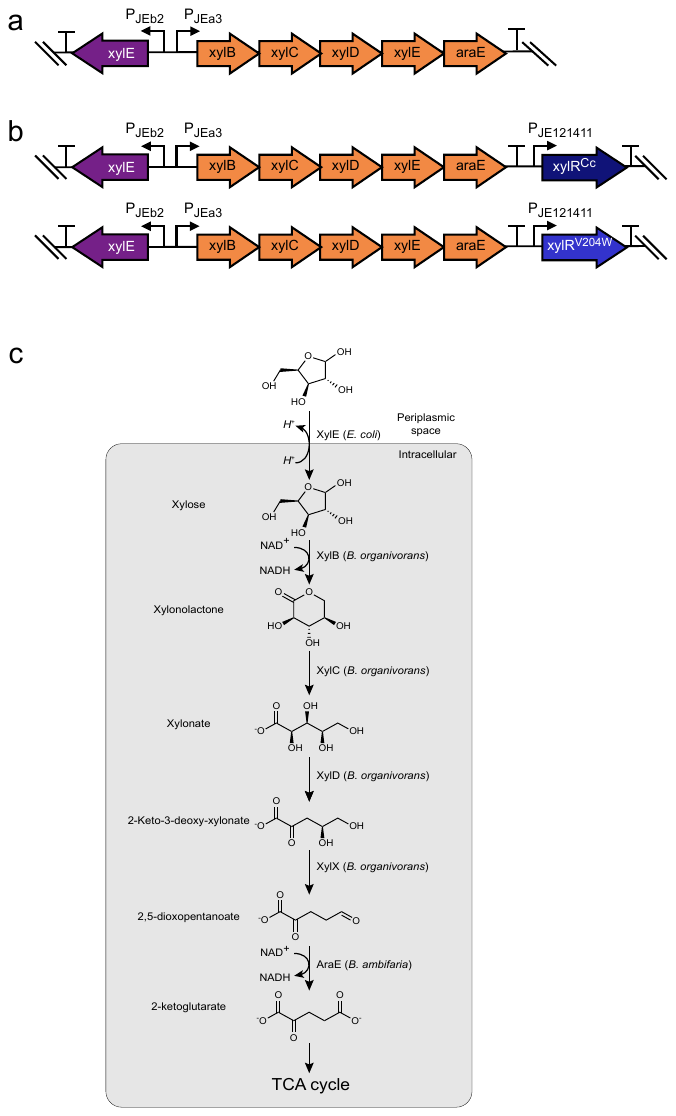

**Supplementary Fig. S1. Oxidative xylose catabolic pathway and expression cassette layout.** **(a)** Arrangement of the static (constitutive) expression oxidative catabolic pathway expression cassette. In purple is the H^+^/xylose symporter from *Escherichia coli*. In orange are the catabolic pathway genes. **(b)** Arrangments of the dynamically regulated catabolic pathway cassettes. Cassettes are identical to the static expression cassette except for the addition of a *C. crescentus* XylR expression cassette appended to the 3’ end of the cassette. **(c)** Diagram of the oxidative xylose catabolic pathway.

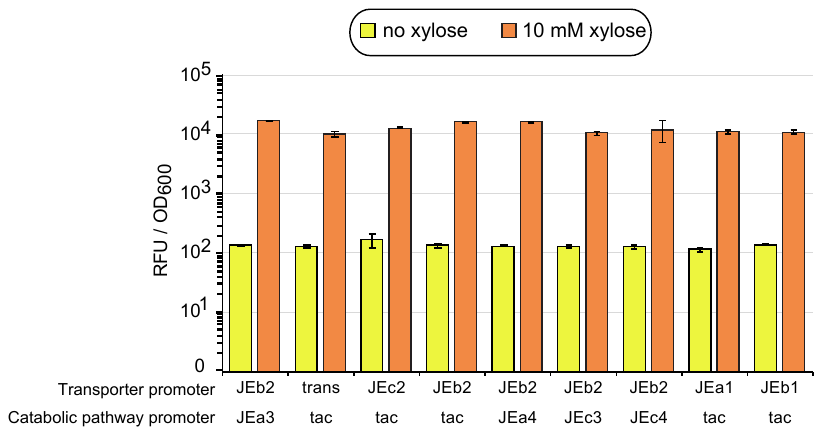

**Supplementary Fig. S2. XylR^Cc^ function is active in strains hosting phosphorylative xylose catabolism cassettes under control of xylose-sensitive promoters.** Graph of mNeonGreen expression in strains hosting constitutive or dynamically regulated phosphorylative xylose catabolism pathway cassettes and a XylR-regulated PJEa3:mNeonGreen cassette integrated into the Bxb1 attP site. Strains were assayed for production of mNeonGreen in defined medium 10 mM glucose and 5 mM xylose as sole carbon sources. Data are presented as the mean values ± standard deviations in three replicates. Source data underlying Supplementary Fig. S2 are provided as a Source Data file.

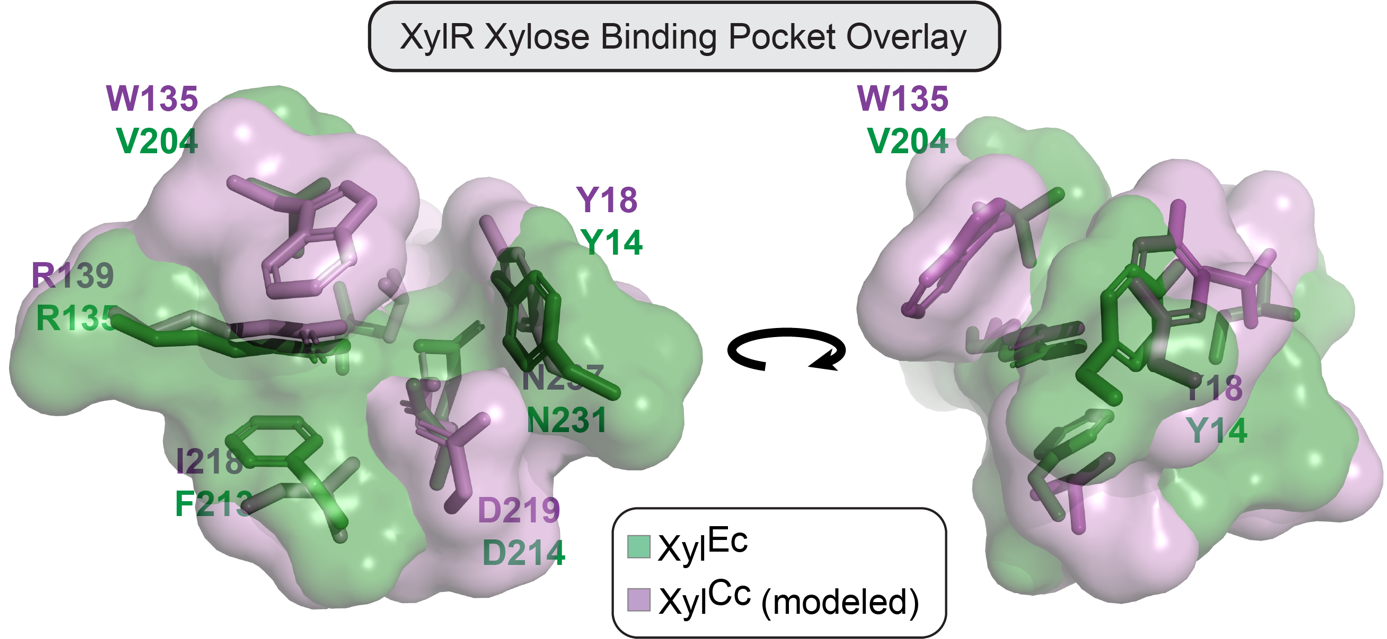

**Supplementary Fig. S3. XylR^Cc^ model structure overlayed with XylR^Ec^ xylose binding pocket.** XylR^Cc^ structure modeled upon PDB ID: 4FE4 is overlayed onto structure 4FE4 of *Escherichia coli* XylR (XylR^Ec^). Overlayed region is rotated 180 degrees around the vertical axis. Images and XylR^Cc^ model structures were generated using PyMol 2.2.3.

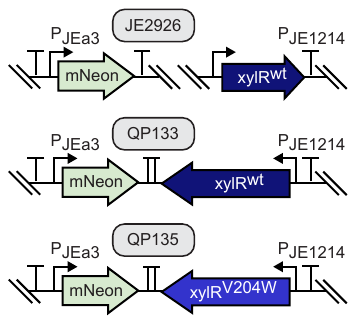

**Supplementary Fig. S4. Diagrams of gene arrangements in strains used to assess XylR^Cc^ and XylR^V204W^ sensitivity to xylose and L-arabinose.** In JE2926 and all strains containing dynamically regulated xylose catabolism cassettes the XylR cassette is distally located from the corresponding mNeonGreen reporter. In QP133 and QP135, used for comparing the sensitivity of wild-type XylR^Cc^ with XylR^V204W^ the two genes are co-located and arranged in a converging arrangement. As such, expression of either gene may be impacted by run-on transcription.

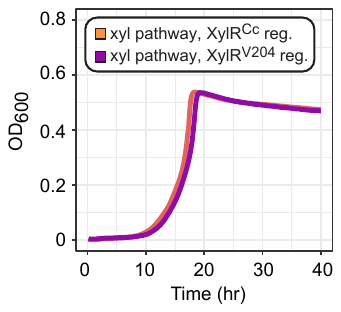

**Supplementary Fig. S5. Growth of non-production strains hosting phosphorylative xylose catabolism cassettes under dynamic control of wild-type or V204W XylR^Cc^ proteins.** Representative growth curves of triplicate cultures in 48-well microtiter plates. Cell density, as measured by OD600, was measured every 10 minutes. Source data underlying Supplementary Fig. S5 are provided as a Source Data file.

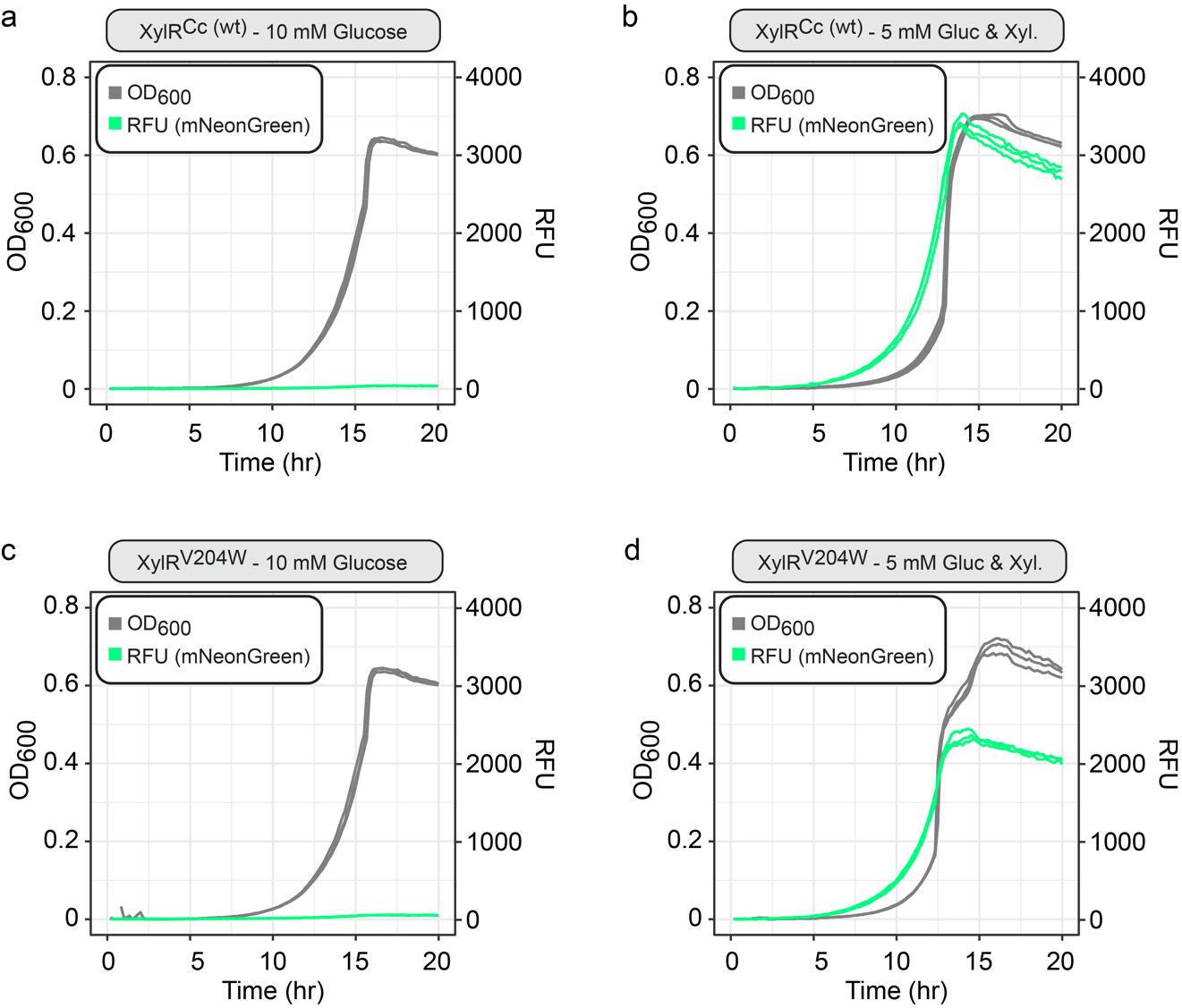

**Supplementary Fig. S6. Growth and production of mNeonGreen by non-production strains hosting dynamically controlled phosphorylative xylose catabolism and mNeonGreen cassettes.** Representative growth curves of triplicate cultures in 96-well microtiter plates. Cell density, as measured by OD600 (gray), and mNeonGreen production, as measured by relative fluourescence – RFU (green), was measured every 10 minutes. Strains utilizing XylR^Cc^ (**a, b**) or XylR^V204W^ (**c, d**) were cultivated with either glucose alone (a,c) or a combination of glucose and xylose **(b,d)**. Source data underlying Supplementary Fig S6 are provided as a Source Data file.

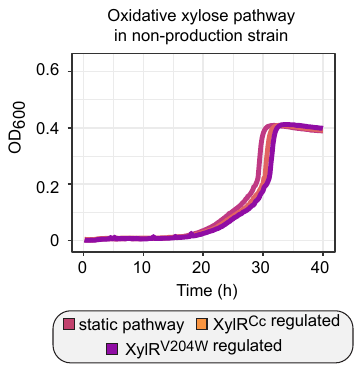

**Supplementary Fig. S7. Growth of non-production strains hosting oxidative xylose catabolism cassettes under static control or dynamic control of wild-type or V204W XylR^Cc^ proteins.** Representative growth curves of triplicate cultures in 48-well microtiter plates. Cell density, as measured by OD600, was measured every 10 minutes. Source data underlying Supplementary Fig. S7 are provided as a Source Data file.

| **Supplementary Table S1. Engineered xylose-sensitive promoters from *Caulobacter crescentus*** | | | |  |
| --- | --- | --- | --- | --- |
| **Source Promoter** | **Promoter Name** | **Source Strain** | **Modification** | **Relative Strength** |
| xylE | JEa1 | *C. crescentus* NA1000 | native | 2.1 |
| xylE | JEa2 | *C. crescentus* NA1000 | optimal -35 | 136.4 |
| xylE | JEa3 | *C. crescentus* NA1000 | optimal -10 | 466.3 |
| xylE | JEa4 | *C. crescentus* NA1000 | optimal -35, optimal -10 | 701.3 |
| xylX | JEb1 | *C. crescentus* NA1000 | native | 1 |
| xylX | JEb2 | *C. crescentus* NA1000 | optimal -35 | 39.8 |
| xylX-K31 | JEc1 | *C. crescentus* K31 | native | 1.6 |
| xylX-K31 | JEc2 | *C. crescentus* K31 | optimal -35 | 71.2 |
| xylX-K31 | JEc3 | *C. crescentus* K31 | optimal -35, improved -10 | 1602.8 |
| xylX-K31 | JEc4 | *C. crescentus* K31 | optimal -35, optimal -10 | 1027.1 |
| --- | tac | synthetic | --- | 917 |

| **Supplementary Table S2. Strains and plasmids used in this work** | |  |
| --- | --- | --- |
| **Name** | **Relevant Genotype** | **Source** |
| *Strains* |  |  |
| NEB 5-alpha F'Iq | *Escherichia coli* F´ *proA^+^B^+^ lacI^q^ ∆(lacZ)M15 zzf::Tn10* (Tet^R^) */ fhuA2∆(argF-lacZ)U169 phoA glnV44 Φ80Δ(lacZ)M15 gyrA96 recA1 relA1 endA1 thi-1 hsdR17* | New England Biolab |
| Epi400 | *Escherichia coli* F^-^ *mcrA Δ(mrr-hsdRMS-mcrBC) Φ80dlacZΔM15 ΔlacX74 recA1 endA1 araD139 Δ(ara, leu)7697 galU galK λ^-^ rpsL (Str^R^) nupG trfA tonA pcnB4 dhfr* | Lucigen |
| JE90 | *Pseudomonas putida* KT2440 *∆hsdR::Bxb1int-attB* | ^1^ |
| JE212 | *P. putida* KT2440 *∆hsdR::Bxb1int-attB ∆gcd* | this work |
| JE1603 | *P. putida* KT2440 *∆hsdR::Bxb1int-attB ∆gcd ∆ampC::P_xylE_-xylEmut* | this work |
| JE2573 | *P. putida* KT2440 *∆hsdR::Bxb1int-attB ∆gcd ∆ampC::P_xylE_-xylEmut ∆aldB-I::XylR^Cc^_v2 (pJE1278)* | this work |
| JE2925 | *P. putida* KT2440 *∆hsdR::Bxb1int-attB ∆gcd ∆ampC::P_xylE_-xylEmut ∆aldB-I::XylR^Cc^_v5 (pJE1338)* | this work |
| JE2926 | *P. putida* KT2440 *∆hsdR::Bxb1int-attB ∆gcd ∆ampC::P_xylE_-xylEmut ∆aldB-I::XylR^Cc^_v6 (pJE1339)* | this work |
| JE2927 | *P. putida* KT2440 *∆hsdR::Bxb1int-attB ∆gcd ∆ampC::P_xylE_-xylEmut ∆aldB-I::XylR^Cc^_v7 (pJE1340)* | this work |
| JE2928 | *P. putida* KT2440 *∆hsdR::Bxb1int-attB ∆gcd ∆ampC::P_xylE_-xylEmut ∆aldB-I::XylR^Cc^_v8 (pJE1341)* | this work |
| JE2597 | *P. putida* KT2440 *∆hsdR::Bxb1int-attL:nptII:P_JEa3_:mNeonGreen:attR ∆gcd* | this work |
| JE3233 | *P. putida* KT2440 *∆hsdR::Bxb1int-attL:nptII:P_JEa3_:mNeonGreen:attR ∆gcd ∆ampC::P_xylE_-xylEmut* | this work |
| JE2606 | *P. putida* KT2440 *∆hsdR::Bxb1int-attL:nptII:P_JEa3_:mNeonGreen:attR ∆gcd ∆ampC::P_xylE_-xylEmut ∆aldB-I::XylR^Cc^_v2* | this work |
| JE2935 | *P. putida* KT2440 *∆hsdR::Bxb1int-attL:nptII:P_JEa3_:mNeonGreen:attR ∆gcd ∆ampC::P_xylE_-xylEmut ∆aldB-I::XylR^Cc^_v5* | this work |
| JE2940 | *P. putida* KT2440 *∆hsdR::Bxb1int-attL:nptII:P_JEa3_:mNeonGreen:attR ∆gcd ∆ampC::P_xylE_-xylEmut ∆aldB-I::XylR^Cc^_v6* | this work |
| JE2945 | *P. putida* KT2440 *∆hsdR::Bxb1int-attL:nptII:P_JEa3_:mNeonGreen:attR ∆gcd ∆ampC::P_xylE_-xylEmut ∆aldB-I::XylR^Cc^_v7* | this work |
| JE2950 | *P. putida* KT2440 *∆hsdR::Bxb1int-attL:nptII:P_JEa3_:mNeonGreen:attR ∆gcd ∆ampC::P_xylE_-xylEmut ∆aldB-I::XylR^Cc^_v8* | this work |
| JE2589 | *P. putida* KT2440 *∆hsdR::Bxb1int-attL:nptII:promoterless_mNeonGreen:attR ∆gcd* | this work |
| JE2590 | *P. putida* KT2440 *∆hsdR::Bxb1int-attL:nptII:P_tac_:mNeonGreen:attR ∆gcd* | this work |
| JE2591 | *P. putida* KT2440 *∆hsdR::Bxb1int-attL:nptII:P_JEb1_:mNeonGreen:attR ∆gcd* | this work |
| JE2592 | *P. putida* KT2440 *∆hsdR::Bxb1int-attL:nptII:P_JEc1_:mNeonGreen:attR ∆gcd* | this work |
| JE2593 | *P. putida* KT2440 *∆hsdR::Bxb1int-attL:nptII:P_JEa1_:mNeonGreen:attR ∆gcd* | this work |
| JE2594 | *P. putida* KT2440 *∆hsdR::Bxb1int-attL:nptII:P_JEc2_:mNeonGreen:attR ∆gcd* | this work |
| JE2595 | *P. putida* KT2440 *∆hsdR::Bxb1int-attL:nptII:P_JEb2_:mNeonGreen:attR ∆gcd* | this work |
| JE2596 | *P. putida* KT2440 *∆hsdR::Bxb1int-attL:nptII:P_JEa2_:mNeonGreen:attR ∆gcd* | this work |
| JE2597 | *P. putida* KT2440 *∆hsdR::Bxb1int-attL:nptII:P_JEa3_:mNeonGreen:attR ∆gcd* | this work |
| QP193 | *P. putida* KT2440 *∆hsdR::Bxb1int-attL:nptII:P_JEa3_:mNeonGreen:xylR^Cc^_­_:attR ∆gcd::P_xylE_:araE ∆ampC::P_xylE_-xylEmut* | this work |
| QP195 | *P. putida* KT2440 *∆hsdR::Bxb1int-attL:nptII:P_JEa3_:mNeonGreen:xylR^V204W^_­_:attR ∆gcd::P_xylE_:araE ∆ampC::P_xylE_-xylEmut* | this work |
| JE3097 | *P. putida* KT2440 *∆hsdR::Bxb1int-attL:nptII:P_JEc3_:mNeonGreen:attR ∆gcd ∆ampC::P_xylE_-xylEmut ∆aldB-I::XylR^Cc^_v6* | this work |
| JE3099 | *P. putida* KT2440 *∆hsdR::Bxb1int-attL:nptII:P_JEa4_:mNeonGreen:attR ∆gcd ∆ampC::P_xylE_-xylEmut ∆aldB-I::XylR^Cc^_v6* | this work |
| JE3101 | *P. putida* KT2440 *∆hsdR::Bxb1int-attL:nptII:P_JEc4_:mNeonGreen:attR ∆gcd ∆ampC::P_xylE_-xylEmut ∆aldB-I::XylR^Cc^_v6* | this work |
| JE4446 | *P. putida* KT2440 *∆hsdR::Bxb1int-attL:nptII:P_JEc3_:mNeonGreen:attR ∆gcd* | this work |
| JE4447 | *P. putida* KT2440 *∆hsdR::Bxb1int-attL:nptII:P_JEa4_:mNeonGreen:attR ∆gcd* | this work |
| JE4448 | *P. putida* KT2440 *∆hsdR::Bxb1int-attL:nptII:P_JEc4_:mNeonGreen:attR ∆gcd* | this work |
| JE2955 | *P. putida* KT2440 *∆hsdR::Bxb1int-attL:nptII:promoterless_mNeonGreen:attR ∆gcd ∆ampC::P_xylE_-xylEmut ∆aldB-I::XylR^Cc^_v6* | this work |
| JE2941 | *P. putida* KT2440 *∆hsdR::Bxb1int-attL:nptII:P_tac_:mNeonGreen:attR ∆gcd ∆ampC::P_xylE_-xylEmut ∆aldB-I::XylR^Cc^_v6* | this work |
| JE2952 | *P. putida* KT2440 *∆hsdR::Bxb1int-attL:nptII:P_JEb1_:mNeonGreen:attR ∆gcd ∆ampC::P_xylE_-xylEmut ∆aldB-I::XylR^Cc^_v6* | this work |
| JE2953 | *P. putida* KT2440 *∆hsdR::Bxb1int-attL:nptII:P_JEc1_:mNeonGreen:attR ∆gcd ∆ampC::P_xylE_-xylEmut ∆aldB-I::XylR^Cc^_v6* | this work |
| JE2954 | *P. putida* KT2440 *∆hsdR::Bxb1int-attL:nptII:P_JEa1_:mNeonGreen:attR ∆gcd ∆ampC::P_xylE_-xylEmut ∆aldB-I::XylR^Cc^_v6* | this work |
| JE2937 | *P. putida* KT2440 *∆hsdR::Bxb1int-attL:nptII:P_JEc2_:mNeonGreen:attR ∆gcd ∆ampC::P_xylE_-xylEmut ∆aldB-I::XylR^Cc^_v6* | this work |
| JE2938 | *P. putida* KT2440 *∆hsdR::Bxb1int-attL:nptII:P_JEb2_:mNeonGreen:attR ∆gcd ∆ampC::P_xylE_-xylEmut ∆aldB-I::XylR^Cc^_v6* | this work |
| JE2939 | *P. putida* KT2440 *∆hsdR::Bxb1int-attL:nptII:P_JEa2_:mNeonGreen:attR ∆gcd ∆ampC::P_xylE_-xylEmut ∆aldB-I::XylR^Cc^_v6* | this work |
| JE3034 | *P. putida* KT2440 *∆hsdR::Bxb1int-attB ∆gcd ∆aldB-I::XylR^Cc^_v6:P_xylE_:xylE:P_tac_:xylAB-talB-tktA (pJE1354)* | this work |
| JE4276 | *P. putida* KT2440 *∆hsdR::Bxb1int-attB ∆gcd ∆aldB-I::XylR^Cc^_v6:P_JEb1_:xylE:P_tac_:xylAB-talB-tktA (pJE1356)* | this work |
| JE4225 | *P. putida* KT2440 *∆hsdR::Bxb1int-attB ∆gcd ∆aldB-I::XylR^Cc^_v6:P_JEa1_:xylE:P_tac_:xylAB-talB-tktA (pJE1357)* | this work |
| JE3189 | *P. putida* KT2440 *∆hsdR::Bxb1int-attB ∆gcd ∆aldB-I::XylR^Cc^_v6:P_JEc2_:xylE:P_tac_:xylAB-talB-tktA (pJE1358)* | this work |
| JE3193 | *P. putida* KT2440 *∆hsdR::Bxb1int-attB ∆gcd ∆aldB-I::XylR^Cc^_v6:P_JEb2_:xylE:P_tac_:xylAB-talB-tktA (pJE1359)* | this work |
| JE3238 | *P. putida* KT2440 *∆hsdR::Bxb1int-attB ∆gcd ∆aldB-I::XylR^Cc^_v6:P_JEb2_:xylE:P_JEa3_:xylAB-talB-tktA (pJE1383)* | this work |
| JE3243 | *P. putida* KT2440 *∆hsdR::Bxb1int-attB ∆gcd ∆aldB-I::XylR^Cc^_v6:P_JEb2_:xylE:P_JEa4_:xylAB-talB-tktA (pJE1384)* | this work |
| JE3244 | *P. putida* KT2440 *∆hsdR::Bxb1int-attB ∆gcd ∆aldB-I::XylR^Cc^_v6:P_JEb2_:xylE:P_JEc3_:xylAB-talB-tktA (pJE1385)* | this work |
| JE3246 | *P. putida* KT2440 *∆hsdR::Bxb1int-attB ∆gcd ∆aldB-I::XylR^Cc^_v6:P_JEb2_:xylE:P_JEc4_:xylAB-talB-tktA (pJE1386)* | this work |
| JE3516 | *P. putida* KT2440 *∆hsdR::Bxb1int-attL:nptII:P_JEa3_:mNeonGreen:attR ∆gcd ∆aldB-I::XylR^Cc^_v6:P_xylE_:xylE:P_tac_:xylAB-talB-tktA (pJE1354)* | this work |
| JE4301 | *P. putida* KT2440 *∆hsdR::Bxb1int-attL:nptII:P_JEa3_:mNeonGreen:attR ∆gcd ∆aldB-I::XylR^Cc^_v6:P_JEb1_:xylE:P_tac_:xylAB-talB-tktA (pJE1356)* | this work |
| JE4300 | *P. putida* KT2440 *∆hsdR::Bxb1int-attL:nptII:P_JEa3_:mNeonGreen:attR ∆gcd ∆aldB-I::XylR^Cc^_v6:P_JEa1_:xylE:P_tac_:xylAB-talB-tktA (pJE1357)* | this work |
| JE3517 | *P. putida* KT2440 *∆hsdR::Bxb1int-attL:nptII:P_JEa3_:mNeonGreen:attR ∆gcd ∆aldB-I::XylR^Cc^_v6:P_JEc2_:xylE:P_tac_:xylAB-talB-tktA (pJE1358)* | this work |
| JE3518 | *P. putida* KT2440 *∆hsdR::Bxb1int-attL:nptII:P_JEa3_:mNeonGreen:attR ∆gcd ∆aldB-I::XylR^Cc^_v6:P_JEb2_:xylE:P_tac_:xylAB-talB-tktA (pJE1359)* | this work |
| JE3375 | *P. putida* KT2440 *∆hsdR::Bxb1int-attL:nptII:P_JEa3_:mNeonGreen:attR ∆gcd ∆aldB-I::XylR^Cc^_v6:P_JEb2_:xylE:P_JEa3_:xylAB-talB-tktA (pJE1383)* | this work |
| JE3519 | *P. putida* KT2440 *∆hsdR::Bxb1int-attL:nptII:P_JEa3_:mNeonGreen:attR ∆gcd ∆aldB-I::XylR^Cc^_v6:P_JEb2_:xylE:P_JEa4_:xylAB-talB-tktA (pJE1384)* | this work |
| JE3520 | *P. putida* KT2440 *∆hsdR::Bxb1int-attL:nptII:P_JEa3_:mNeonGreen:attR ∆gcd ∆aldB-I::XylR^Cc^_v6:P_JEb2_:xylE:P_JEc3_:xylAB-talB-tktA (pJE1385)* | this work |
| JE3521 | *P. putida* KT2440 *∆hsdR::Bxb1int-attL:nptII:P_JEa3_:mNeonGreen:attR ∆gcd ∆aldB-I::XylR^Cc^_v6:P_JEb2_:xylE:P_JEc4_:xylAB-talB-tktA (pJE1386)* | this work |
| JE4280 | *P. putida* KT2440 *∆hsdR::Bxb1int-attB ∆gcd ∆aldB-I::XylR^V204W^_v6:P_JEb2_:xylE:P_JEa3_:xylAB-talB-tktA (pJE1517)* | this work |
| JE4220 | *P. putida* KT2440 *∆hsdR::Bxb1int-attB ∆gcd ∆aldB-I::P_JEb2_:xylE:P_JEa3_:xylDXBC (pJE1503)* | this work |
| JE4284 | *P. putida* KT2440 *∆hsdR::Bxb1int-attB ∆gcd ∆aldB-I::XylR^Cc^_v6:P_JEb2_:xylE:P_JEa3_:xylDXBC (pJE1504)* | this work |
| JE4286 | *P. putida* KT2440 *∆hsdR::Bxb1int-attB ∆gcd ∆aldB-I::XylR^V204W^_v6:P_JEb2_:xylE:P_JEa3_:xylDXBC (pJE1518)* | this work |
| JE4302 | *P. putida* KT2440 *∆hsdR::Bxb1int-attL:nptII:P_JEa3_:mNeonGreen:attR ∆gcd ∆aldB-I::XylR^V204W^_v6:P_JEb2_:xylE:P_JEa3_:xylAB-talB-tktA (pJE1517)* | this work |
| JE4299 | *P. putida* KT2440 *∆hsdR::Bxb1int-attL:nptII:P_JEa3_:mNeonGreen:attR ∆gcd ∆aldB-I::P_JEb2_:xylE:P_JEa3_:xylDXBC (pJE1503)* | this work |
| JE4303 | *P. putida* KT2440 *∆hsdR::Bxb1int-attL:nptII:P_JEa3_:mNeonGreen:attR ∆gcd ∆aldB-I::XylR^Cc^_v6:P_JEb2_:xylE:P_JEa3_:xylDXBC (pJE1504)* | this work |
| JE4304 | *P. putida* KT2440 *∆hsdR::Bxb1int-attL:nptII:P_JEa3_:mNeonGreen:attR ∆gcd ∆aldB-I::XylR^V204W^_v6:P_JEb2_:xylE:P_JEa3_:xylDXBC (pJE1518)* | this work |
| CJ442 | *P. putida* KT2440 *ΔcatRBCA::P_tac_:catA ΔpcaHG::P_tac_:aroY:ecdB:asbF ΔpykA::aroG-D146N:aroY:ecdB:asbF ΔpykF Δppc Δpgi-1 Δpgi-2* | ^2^ |
| JE2985 | *P. putida* KT2440 *ΔcatRBCA::P_tac_:catA ΔpcaHG::P_tac_:aroY:ecdB:asbF ΔpykA::aroG-D146N:aroY:ecdB:asbF ΔpykF Δppc Δpgi-1 Δpgi-2 ∆hsdR::Bxb1int-attB* | this work |
| JE4346 | *P. putida* JE2985 *∆aldB-I::P_JEb2_:xylE:P_JEa3_:xylDXBC (pJE1503)* | this work |
| JE4349 | *P. putida* JE2985 *∆aldB-I::XylR^Cc^_v6:P_JEb2_:xylE:P_JEa3_:xylDXBC (pJE1504)* | this work |
| *Plasmids* |  |  |
| pK18mobsacB | pUC origin, *nptII*, *sacB*, *P_lac_:mcs-lacZa* | ^3^ |
| pJE365 | pK18mobsacB *∆gcd* cassette | ^4^ |
| pJE382 | pUC origin, *nptII*, *sacB*, *mcs-lacZa* (promoterless multiple cloning site, based on pK18mobsacB) | ^4^ |
| pJE443 | pK18mobsacB *∆ampC::P_xylE_-xylE:P_tac_-xylAB:talB:tktA* cassette | ^4^ |
| pΔ*hsdR*::BxB1int-attB | pK18mobsacB-derivative for replacement of PP_4740 (*hsdR*) with codon-optimized BxB1int-attB | ^1^ |
| pJE990 | pUC origin, *nptII*, *mNeonGreen* (promoterless), *Bxb1 attP* | ^1^ |
| pJE1096 | pJE990 P_tac_:*mNeonGreen* | this work |
| pJE1278 | pK18mobsacB *∆aldB-I::XylR^Cc^_v2 (native XylR promoter/RBS)* | this work |
| pJE1279 | pJE990 P_JEb1_:*mNeonGreen* | this work |
| pJE1280 | pJE990 P_JEc1_:*mNeonGreen* | this work |
| pJE1281 | pJE990 P_JEa1_:*mNeonGreen* | this work |
| pJE1282 | pJE990 P_JEc2_:*mNeonGreen* | this work |
| pJE1283 | pJE990 P_JEb2_:*mNeonGreen* | this work |
| pJE1284 | pJE990 P_JEa2_:*mNeonGreen* | this work |
| pJE1285 | pJE990 P_JEc3_:*mNeonGreen* | this work |
| pJE1286 | pJE990 P_JEa3_:*mNeonGreen* | this work |
| pJE1338 | pK18mobsacB *∆aldB-I::XylR^Cc^_v5 (JE151111 promoter, JER08 RBS)* | this work |
| pJE1339 | pK18mobsacB *∆aldB-I::XylR^Cc^_v6 (JE121411 promoter, pJE483 RBS)* | this work |
| pJE1340 | pK18mobsacB *∆aldB-I::XylR^Cc^_v7 (tac promoter, JER08 RBS)* | this work |
| pJE1341 | pK18mobsacB *∆aldB-I::XylR^Cc^_v8 (tac promoter, pJE483 RBS)* | this work |
| pJE1354 | pK18mobsacB *∆aldB-I::XylR^Cc^_v6:P_xylE_:xylE:P_tac_:xylAB-talB-tktA* | this work |
| pJE1356 | pK18mobsacB *∆aldB-I::XylR^Cc^_v6:P_JEb1_:xylE:P_tac_:xylAB-talB-tktA* | this work |
| pJE1357 | pK18mobsacB *∆aldB-I::XylR^Cc^_v6:P_JEa1_:xylE:P_tac_:xylAB-talB-tktA* | this work |
| pJE1358 | pK18mobsacB *∆aldB-I::XylR^Cc^_v6:P_JEc2_:xylE:P_tac_:xylAB-talB-tktA* | this work |
| pJE1359 | pK18mobsacB *∆aldB-I::XylR^Cc^_v6:P_JEb2_:xylE:P_tac_:xylAB-talB-tktA* | this work |
| pJE1366 | pJE990 P_JEa4_:*mNeonGreen* | this work |
| pJE1367 | pJE990 P_JEc4_:*mNeonGreen* | this work |
| pJE1383 | pK18mobsacB *∆aldB-I::XylR^Cc^_v6:P_JEb2_:xylE:P_JEa3_:xylAB-talB-tktA* | this work |
| pJE1384 | pK18mobsacB *∆aldB-I::XylR^Cc^_v6:P_JEb2_:xylE:P_JEa4_:xylAB-talB-tktA* | this work |
| pJE1385 | pK18mobsacB *∆aldB-I::XylR^Cc^_v6:P_JEb2_:xylE:P_JEc3_:xylAB-talB-tktA* | this work |
| pJE1386 | pK18mobsacB *∆aldB-I::XylR^Cc^_v6:P_JEb2_:xylE:P_JEc4_:xylAB-talB-tktA* | this work |
| pJE1503 | pK18mobsacB *∆aldB-I::P_JEb2_:xylE:P_JEa3_:xylDXBC* | this work |
| pJE1504 | pK18mobsacB *∆aldB-I::XylR^Cc^_v6:P_JEb2_:xylE:P_JEa3_:xylDXBC* | this work |
| pJE1517 | pK18mobsacB *∆aldB-I::XylR^V204W^_v6:P_JEb2_:xylE:P_JEa3_:xylAB-talB-tktA* | this work |
| pJE1518 | pK18mobsacB *∆aldB-I::XylR^V204W^_v6:P_JEb2_:xylE:P_JEa3_:xylDXBC* | this work |
| pJE942 | pK18mobsacB *∆ampC::P_xylE_-xylEmut* | this work |
| pQP133 | pJE990 *P_JEa3_:mNeonGreen:xylR^Cc^* | this work |
| pQP135 | pJE990 *P_JEa3_:mNeonGreen:xylR^V204W^* | this work |

| Supplemental Table S3. DNA oligos used in this work | | |
| --- | --- | --- |
| **Oligo Name** | **Sequence** (5'-3') | **Purpose** |
| oJE93 | GGCGTTGCTGGAAGAGTATT | flanking primers to screen for replacement of *ampC* with xylEmut cassette |
| oJE94 | ACCACTGCCAGCAGAATTG | flanking primers to screen for replacement of *ampC* with xylEmut cassette |
| oJE461 | AGCCAGCTGTTCTTGTCCAT | screen for presence of xylEmut |
| oJE462 | CCATGTACATTGCCGAACTG | screen for presence of xylEmut |
| oJE66 | catgtagttgtaggcgtcttc | screen for insertion of pJE990-based plasmids into Bxb1-attB site |
| oJE536 | aaaaccgcccagtctagctatcg | screen for insertion of pJE990-based plasmids into Bxb1-attB site |
| oJE446 | CGGGTACAGGTGGTAGCAGT | aldB-I internal primers to screen for replacement of *aldB-I* with various constructs |
| oJE447 | CACCGTACTGCTCGAAGTCA | aldB-I internal primers to screen for replacement of *aldB-I* with various constructs |
| oJE546 | gctgttgccatcgatcagt | ampC internal primers to ensure gene deletion |
| oJE547 | acgaccagttacaggccaag | ampC internal primers to ensure gene deletion |
| oJE1129 | CCCAGATCGTGATCGAAAAG | xylRCc internal primers |
| oJE1130 | GAAATAGCCCTGCTCCACAC | xylRCc internal primers |
| oJE175 | CGACCAGCGGGTTACC | aldB-I flanking primers to screen for replacement of *aldB-I* with various constructs |
| oJE176 | CTGAACATTTCGTCATGGCTG | aldB-I flanking primers to screen for replacement of *aldB-I* with various constructs |
| oJE1322 | CTTTTAACCGTTCGACAGC | hsdR flanking primers to screen for insertion of Bxb1 integrase and attB |
| oJE1323 | CATCAAGACCAACCTGCTG | hsdR flanking primers to screen for insertion of Bxb1 integrase and attB |
| oJE1324 | CGAAATCTACCTGTCGCTC | hsdR internal primers to screen for deletion of hsdR |
| oJE1325 | CGTGATTGATATCGTCGC | hsdR internal primers to screen for deletion of hsdR |
